# Adjuvant Discovery via a High Throughput Screen using Human Primary Mononuclear Cells

**DOI:** 10.1101/2022.06.17.496630

**Authors:** Katherine Chew, Branden Lee, Simon D. van Haren, Etsuro Nanishi, Timothy O’Meara, Jennifer B. Splaine, Maria DeLeon, Dheeraj Soni, Hyuk-Soo Seo, Sirano Dhe-Paganon, Al Ozonoff, Jennifer A. Smith, Ofer Levy, David J. Dowling

## Abstract

We describe a methodology utilizing high-throughput and high-content screening for novel adjuvant candidates that was used to screen a library of ∼2,500 small molecules via a 384-well quantitative combined cytokine and flow cytometry immunoassay in primary human peripheral blood mononuclear cells (PBMCs) from 4 healthy adult study participants. Hits were identified based on their induction of soluble cytokine (TNF, IFNγ and IL-10) secretion and PBMC maturation (CD 80/86, Ox40, and HLA-DR) in at least two of the four donors screened. From an initial set of 197 compounds identified using these biomarkers—an 8.6% hit rate—we downselected to five scaffolds that demonstrated robust efficacy and potency *in vitro* and evaluated the top hit, vinblastine sulfate, for adjuvanticity *in vivo*. Vinblastine sulfate significantly enhanced murine humoral responses to recombinant SARS-CoV-2 spike protein, including a four-fold enhancement of IgG titer production when compared to treatment with the spike antigen alone. Overall, we outline a methodology for discovering immunomodulators with adjuvant potential via high-throughput screening of PBMCs *in vitro* that yielded a lead compound with *in vivo* adjuvanticity.

**Motivation:** Vaccines are a key biomedical intervention to prevent the spread of infectious diseases, but their efficacy can be limited by insufficient immunogenicity and nonuniform reactogenic profiles. Adjuvants are molecules that potentiate vaccine responses by inducing innate immune activation. However, there are a limited number of adjuvants in approved vaccines, and current approaches for preclinical adjuvant discovery and development are inefficient. To enhance adjuvant identification, we developed a protocol based on *in vitro* screening of human primary leukocytes.

## Introduction

Other than clean drinking water, vaccines are the most impactful public health intervention in history. The World Health Organization estimates that childhood vaccines have eliminated diseases like measles and rubella from >80 countries and prevent ∼2.5 million deaths each year (MacDonald et al., 2020). In the United States, the economic burden of vaccine-preventable diseases was $9 billion in 2015 alone (Ozawa et al., 2016). However, the recent emergence of the severe acute respiratory syndrome coronavirus 2 (SARS-CoV-2) has precipitated a global pandemic, which has resulted in >500 million cases and nearly six million deaths as of April 2022. Early estimates anticipate nearly ∼$16 trillion in economic losses from this pandemic (Cutler and Summers, 2020). Overall, there remains an unmet need for novel vaccines to protect against microbes for which there are no currently approved immunizations (e.g., respiratory syncitial virus, human immunodeficiency virus) or for which improved vaccines are needed (e.g., pertussis, tuberculosis, influenza, coronavirus).

Several challenges exist in the modern vaccine development, including waning immunity and the need to vaccinate immunologically distinct populations such as older adults. The protection conferred by vaccines for a range of infections such as influenza, mumps, and pertussis wanes over time (Cohen, 2019). Waning immunity raises a deeper challenge, considering the prevalence of antigenic drift common in the influenza virus (Collier et al., 2021) and the high mutability of certain pathogens, such as emerging variants of SARS-CoV-2 (Lauring and Malani, 2021). Even vaccines capable of conferring broad protection against these heterovariant strains may not be sufficiently effective in individuals with distinct immunity, such as the immunocompromised, neonates/infants, and older adults (Angelidou et al., 2020; Collier *et al*., 2021; Pettengill et al., 2016; van Haren et al., 2016b)

These challenges highlight the need for novel adjuvants that enhance the immunogenicity and effectiveness of vaccines (Nanishi et al., 2020; O’Hagan and Valiante, 2003; Pulendran et al., 2021; Reed et al., 2013). Adjuvants represent a broad class of vaccine components that improve vaccine efficacy, stability, and/or durability. Of these broad class of adjuvants, a subset of small molecules function to improve the immunogenicity of vaccines by activating toll-like receptors (TLRs) on antigen presenting cells (APCs), such as dendritic cells or macrophages (Pulendran *et al*., 2021). This activity stimulates innate immune responses, including the induction of proinflammatory cytokines. This innate immune activation then helps promote the efficacy and durability of the adaptive immune responses, from improved antibody (Ab) functionality and magnitude to enhanced CD4^+^ T helper cell responses (Burny et al., 2017; Francica et al., 2017; Petitdemange et al., 2019).

The most longstanding adjuvants are aluminum salts (alum), which induce a robust humoral response in a safe and cost-effective manner (Kool et al., 2012). However, alum has limited ability to elicit strong T helper 1 (Th1) cellular immunity, instead skewing largely toward a Th2 response, especially in a pediatric context (Dowling and Levy, 2015). This skew limits the effectiveness and applicability of alum as an adjuvant in vaccines against intracellular pathogens (Oleszycka et al., 2018). These limitations have precipitated the development of novel adjuvant systems, including squalene-in-water-emulsions, monophosphoryl lipid A (MPLA), and other small molecule combinations (Reed *et al*., 2013). Growing the adjuvant pipeline by providing a consortium of immunostimulatory profiles will enable development of more precise, adaptable, and effective vaccine formulations (Soni et al., 2020b) (NIAID, 2018).

High-throughput screening (HTS) is a powerful approach for the discovery of new adjuvants. HTS enables rapid and effective investigation of the immunomodulatory profiles of thousands of small molecules to identify potential adjuvanticity. Most successful screening campaigns leverage well-defined cell lines to discover immunomodulatory qualities using very precise but limited measurements (Hu et al., 2021; Spangenberg et al., 2021; Wong et al., 2015). While these screens yield precise and controlled results, they do not always accurately capture the fuller diversity of the immune system since these non-diverse cell lines cannot recapitulate the full human immune response. Human primary immune cells may more accurately model the diverse human immune system by enabling the simultaneous study of a variety of relevant cell types. However, several challenges have been identified in screening paradigms that leverage primary cells (Dunne et al., 2009), contributing to the dearth of primary cell screening campaigns.

Innate immune activation is important to robust responses across a range of vaccines (Fourati et al., 2021), providing a conceptual basis for an adjuvant screen based on biomarkers of innate immune activation. *In vitro* systems that leverage human leukocytes can accurately model and have predicted the action of adjuvants and vaccines *in vivo* (Levy et al., 2006; Nanishi et al., 2022; Oh et al., 2016; Philbin et al., 2012; Sanchez-Schmitz et al., 2018). Similarly, the utilization of autologous plasma, which contains age-specific soluble mediators of immunity, may help capture relevant leukocytic responses (England et al., 2021; Pettengill et al., 2014). Here, we describe a robust methodology for HTS utilizing human PBMCs cultured in autologous plasma for multiplexed soluble (TNF, IFNγ and IL-10) and cellular (CD80/86, Ox40, and HLA-DR) biomarkers, through which we identified four leading hits that were validated and titratable *in vitro*, with two of these compounds demonstrating adjuvanticity *in vivo*.

## Results

### Initial Primary Cell Screen Identifies 197 Hits

To discover novel adjuvant candidates, we screened 2,296 compounds from seven distinct library plates to assess their potential immunostimulatory qualities. We stimulated cryopreserved human PBMCs from four young adult study participants (aged 23–27) with these compounds for 72 hours (Figure 1A-B). To capture the diversity of immunological profiles and activity, we used a multiplexed readout platform that captured cellular receptor activity of B cells, T cells, and monocytes using flow cytometry. We also measured cytokine production (TNF, IFN-γ, and IL-10) using AlphaLISA technology (Figure 1C). This multi-analyte system captures diverse dimensions of soluble cytokine/interferon primary innate immune responses while also enabling the discovery of cellular subtype activation and maturation. This comprehensive soluble and cellular approach is particularly attractive since effective adjuvants activate both innate and adaptive immunity (Coffman et al., 2010; Sui et al., 2010).

**Figure 1:**
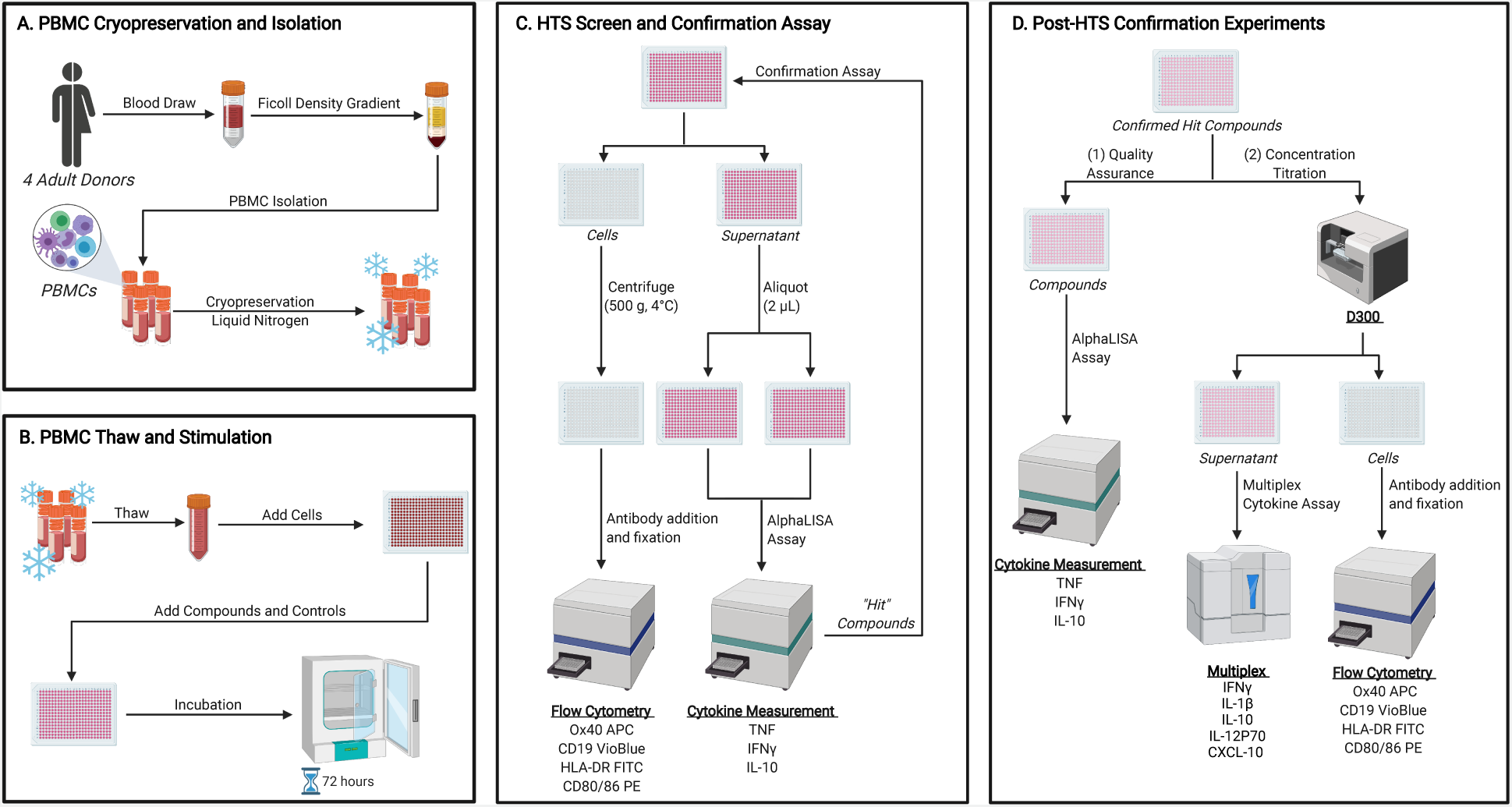
Graphic overview of a multiplexed high throughput screen for novel immunomodulators using human primary cells. **A**. Human peripheral blood mononuclear cells (PBMCs) were isolated and cryopreserved in 10% DMSO and 90% autologous plasma for long-term liquid nitrogen storage. **B**. The cells were thawed, cultured in 10% autologous plasma and 90% DMEM, and distributed onto a 384 well plate. These wells stimulated with screening compounds dissolved in DMSO for a 72 hour incubation at 37°C. **C**. PBMCs were fixated with antibodies for iQue advanced flow cytometry, and cytokine concentrations measured in supernatants employing AlphaLISA. Hit compounds from this initial screen were selected for confirmation assay. **D**. Confirmed hits were tested in a quality assurance (QA) experiment by AlphaLISA and further tested in a dose titration and multiplexed cytokine assay.

Using a compound concentration of 33 µM, hits were identified as activating zero (black), one (yellow), two (green), three (blue) or four (red) participant samples, with Z scores ranging from ∼2 – 30 for cytokine induction (Figure 2A), ∼2 – 4 for monocyte activation/maturation (Figure 2B), ∼2 – 60 for T-cell activation (Figure 2C), ∼2 – 15 for B-cell activation/maturation (Figure 2D). This screen initially identified 197 compounds that demonstrated a sufficient immunomodulatory characteristic in any of the readouts for ≥ 2 of the 4 study participants, providing a hit rate of 8.6% (Figure 2E). Using TNF as an example readout, 89 compounds hit on one donor sample (yellow), 21 hit on two (green), six hit on three (blue) and four hit on four (red) (Figure 2A). When considering both receptor-based and cytokine readouts, these 197 compounds demonstrated non-uniform and variable immunological activity (Figure 2A-E). Similarly, 65 compounds (2.8%) of these 197 (8.6%) were active in ≥3 of the 4 study participants and only 15 (0.65%) were active in all 4 participants (Figure 2E). The strong negative correlation (R^2^ = 0.9958) between number of study participants and a natural log transformed potential hit rate explains a negative, exponential relationship between these two factors. There were exponentially fewer hit compounds as sample size increased, reflecting the diversity of immune capabilities and responsiveness represented in the human population (Tsuchiya and Ohashi, 2015).

**Figure 2:**
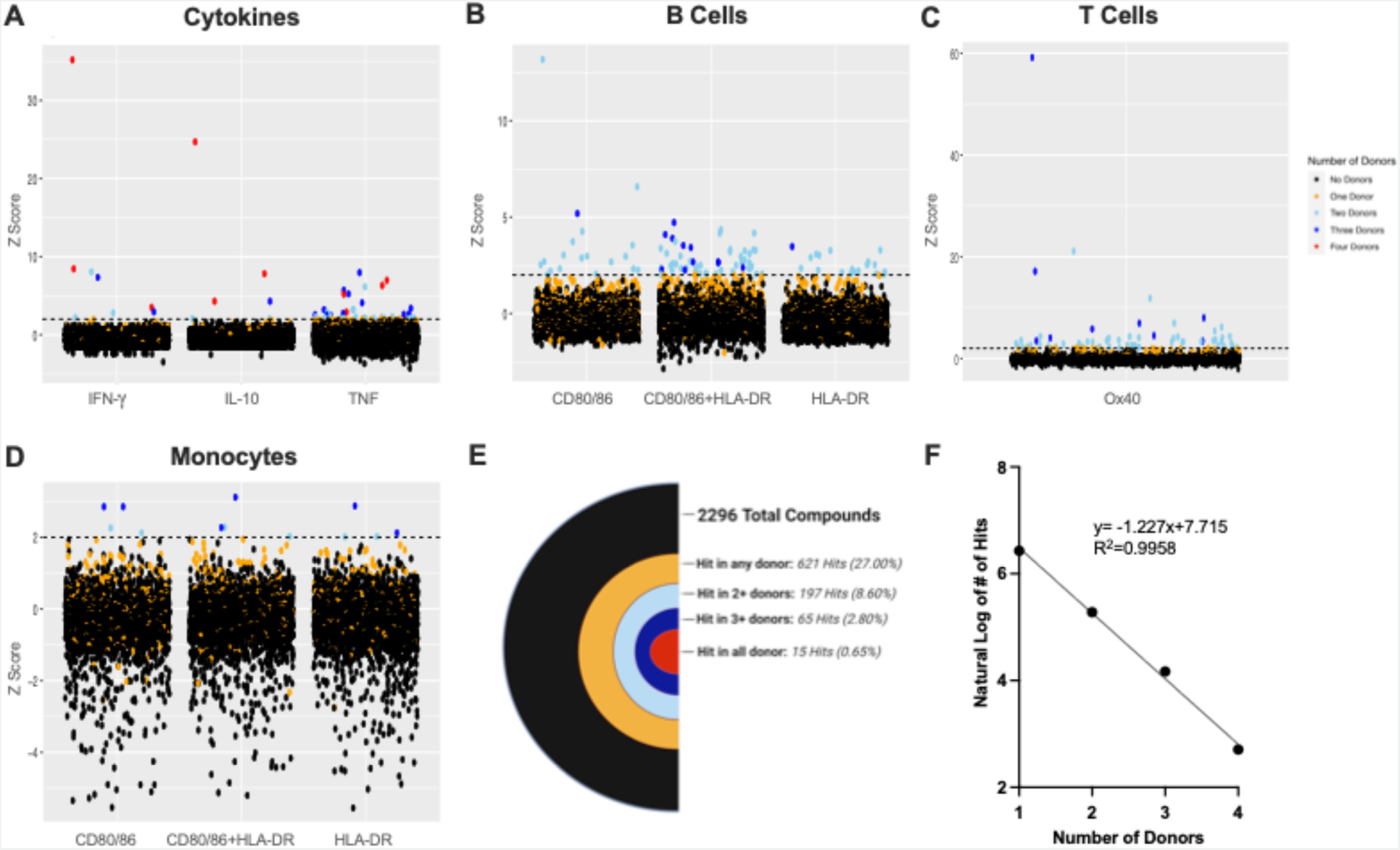
Overview of primary High Throughput Screen data. **A**. Distribution of z scores, indicating induction of the cytokines (IFN, IL-10, and TNF). **B**. Distribution of z scores, representing the percentage of monocytes with activated HLA-DR and/or CD80/86 receptors. **C**. Distribution of z scores, representing the percentage of T-Cells with activated Ox40 receptors. **D**. Distribution of z scores, representing the percentage of B-Cells with activated HLA-DR and/or CD80/86 receptors. **E.** graphical representation of small molecule hit rate in varying number of donors. natural log of hit rate. **F**. Strong negative correlation between number of donors. Each dot represents one of the 2,296 compounds from the primary screen. The dashed line represents the donor threshold of a z score of 2 and representative z scores were chosen such that any compound above the threshold represents a hit in the screen.

Typically, biological variability is substantially greater than technical variability (Blainey et al., 2014). For the same reason of variability within the population, a demonstration of consistency and reproducibility of the results is imperative. To this end, we tested each compound twice and measured absolute value differences between technical replicates (i.e., if compound X induces Y (replicate 1) and Z (replicate 2) pg/ml TNF, the variability will be calculated at |Y-Z|). Across all study participants, compound-induced biomarkers demonstrated a high level of reproducibility, with most absolute value differences between identical wells aggregating near zero (Figures 3A-D). Of the readouts measured, IFNγ and IL-10 (Figures 3B-C) demonstrated the greatest reproducibility between replicates, followed by TNF (Figure 3 A), while combined flow cytometry demonstrates the least (Figure 3D). Similarly, across all assay plates, most positive (R848) and negative (DMSO) control values present coefficient of variance values < 50 (Figure 3E). Even if the hit-calling methodology adjusts and accounts for outliers, these data indicate that the results are highly consistent and reliable across all readout systems.

**Figure 3:**
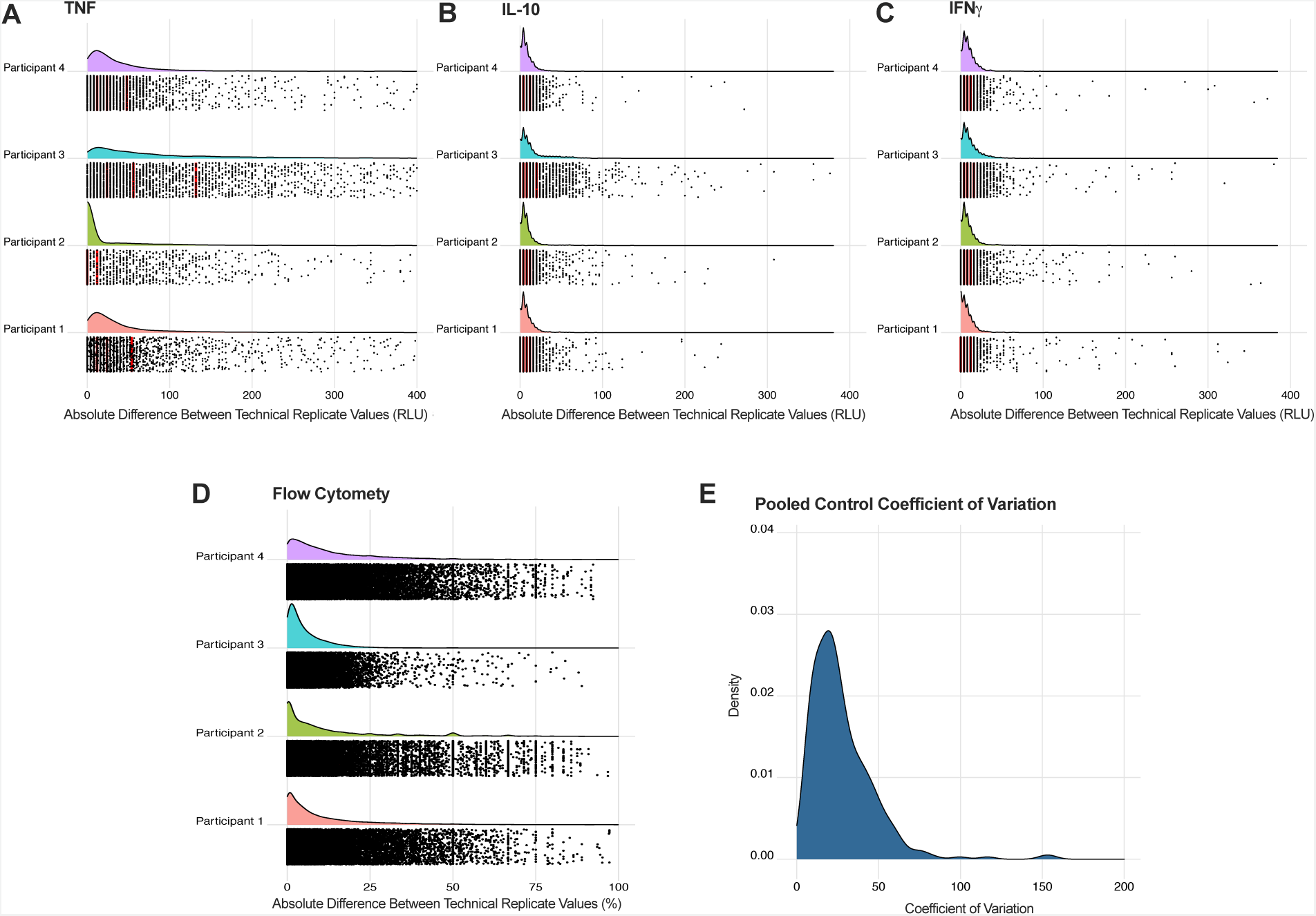
Consistency of HTS results across study participants. **A-D**. Distribution of absolute value differences between technical replicate values for cytokine (TNF, IL-10, and IFN), and high-throughput flow cytometry results (CD80/86, HLA-DR. OX-40), respectively. Each dot represents an individual absolute value difference between technical replicates. Red lines indicate the quantile lines for the absolute value differences for each donor. **E**. Distribution of the coefficient of variation (CV) for positive control compounds (R848 and CpG ODN) across all study participants. Dotted lines represent thresholds of CV quality measures. Dashed lines demarkate typically accepted thresholds for excellent (<10), good (<30), acceptable (<50), and poor (>50) CV values.

### Hit Confirmation Assays Reconfirm 19 Compounds from Screen

To validate the potential hits attained from the initial screen, we re-tested the 197 potential hit compounds using the same quantitative assays as the initial screen. Results from this confirmation screen informed the downselection of 20 compounds for their ability to induce a Z score >2 for cytokine production (Figure 4A) and/or expression of both monocytic cell-surface antigens (Figure 4B). Bearing in mind the key role of monocytes in regulating both innate and adaptive immunity (Leon and Ardavin, 2008; Varga and Foell, 2018), monocyte receptor expression was prioritized over the other flow cytometry-based measurements. To capture this dual function, compounds were evaluated for their ability to induce expression of the cell-surface antigens CD80/86 and HLA-DR. These complexes play an immunostimulatory role in T-cell activation (Boussiotis et al., 1993; Cheadle, 1993; Fleischer et al., 1996; Wang et al., 2018) and can skew the immune system towards Th1 or Th2 differentiation (Slavik et al., 1999), suggesting potential clinical relevance for adjuvanticity (Dhar et al., 2003; Martins et al., 2015) and a possible association with APC maturation (Dowling et al., 2008). As a result, only these 20 compounds that demonstrated the ability to cause sufficient induction of both CD80/86 and HLA-DR expression (Z score >2) were selected for further screening. Confirmation of one biomarker did not necessarily correspond to confirmation of other biomarkers (Supplementary Figure 1). Further, not all hits in the primary screen were validated in the confirmation assays, likely due to the more stringent validation parameters. With each successive confirmation or validation experiment, compounds were downselected with increasingly rigorous standards.

**Figure 4:**
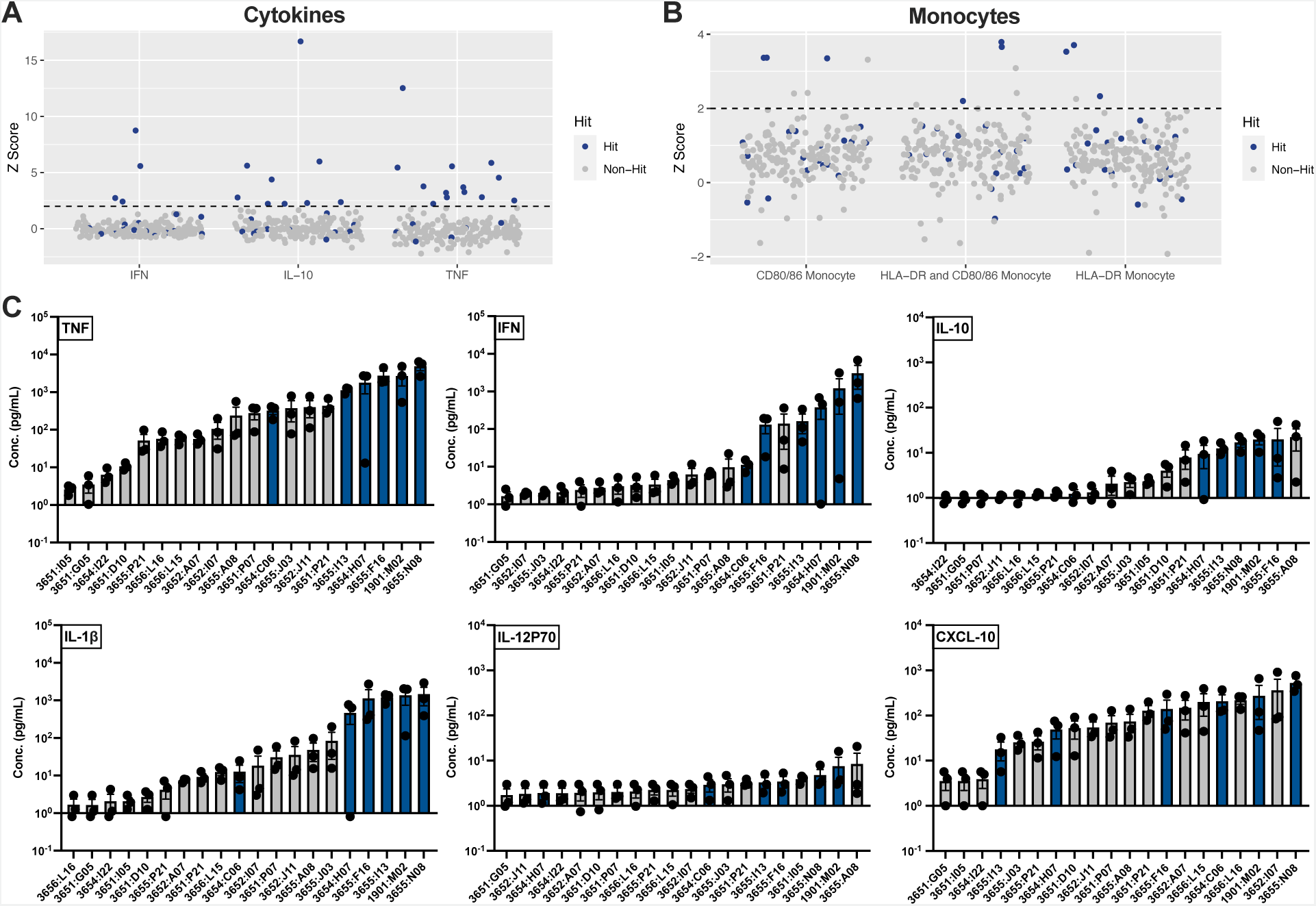
Confirmation and titration of high throughput screening hits. **A**. Distribution of z scores, representing the percentage of monocytes with activated HLA-DR and/or CD80/86 receptors. The dashed line represents the hit threshold of a z score of 2. All receptor combinations must be increased for a compound to be classified as a confirmed hit. **B**. Distribution of z scores, representing production of IFN, IL-10, and TNF. The dashed line represents the hit threshold of a z score of 2. **C**. Rank order of mean cytokine induction (± SEM) at 11 µM for the 19 compounds selected for the multiplexed concentration titration assay based on the results of the monocyte receptor activation and cytokine induction in the confirmation experiment.

### Concentration Titration and Cytokine QA Assays Identify Four Lead Hits

The 19 compounds validated in the hit confirmation experiment were next re-arrayed onto separate plates from their original chemical library plates, and tested at concentrations ranging from 0.25 – 33 µM. For this portion of the screen, we carefully chose a panel of six cytokines, three as a confirmation from the primary screen and three associated with DC maturation and Th-polarization (Arango Duque and Descoteaux, 2014; Striz et al., 2014). Initially, at the therapeutically relevant dosage of 11 µM, four compounds (Figure 4C, Blue) were identified as lead candidates due to consistent performances across these cytokine biomarkers.

These four lead compounds were lexibulin, amphotericin B, silmitasertib, and vinblastine sulfate (Figure 5). At maximal efficacy, all four candidates demonstrated a clear separation from the other compounds (Figure 4C). Additionally, these compounds demonstrated an important concentration-dependent response for many of the cytokines measured (Figure 5). As a result of this demonstrated efficacy, titratability, and potency, these four compounds were chosen as lead adjuvant candidates. The selection criteria prioritized efficacy and titratability while maintaining a lower floor for potency (Supplementary Table 2). This decision accounted for the importance of potency, as measured by Log EC_50_, in ensuring any true adjuvants could be administered at therapeutically relevant dosages while also aiming to find compounds that could potentially induce a clinically relevant response. Overall, TNF was the most effectively induced cytokine, with lexibulin demonstrating the largest maximal efficacy of the downselected candidates (3867 pg/mL), followed by silmitasertib (1627 pg/mL), vinblastine (1622 pg/mL), and lastly amphotericin B (844.8 pg/mL).

**Figure 5:**
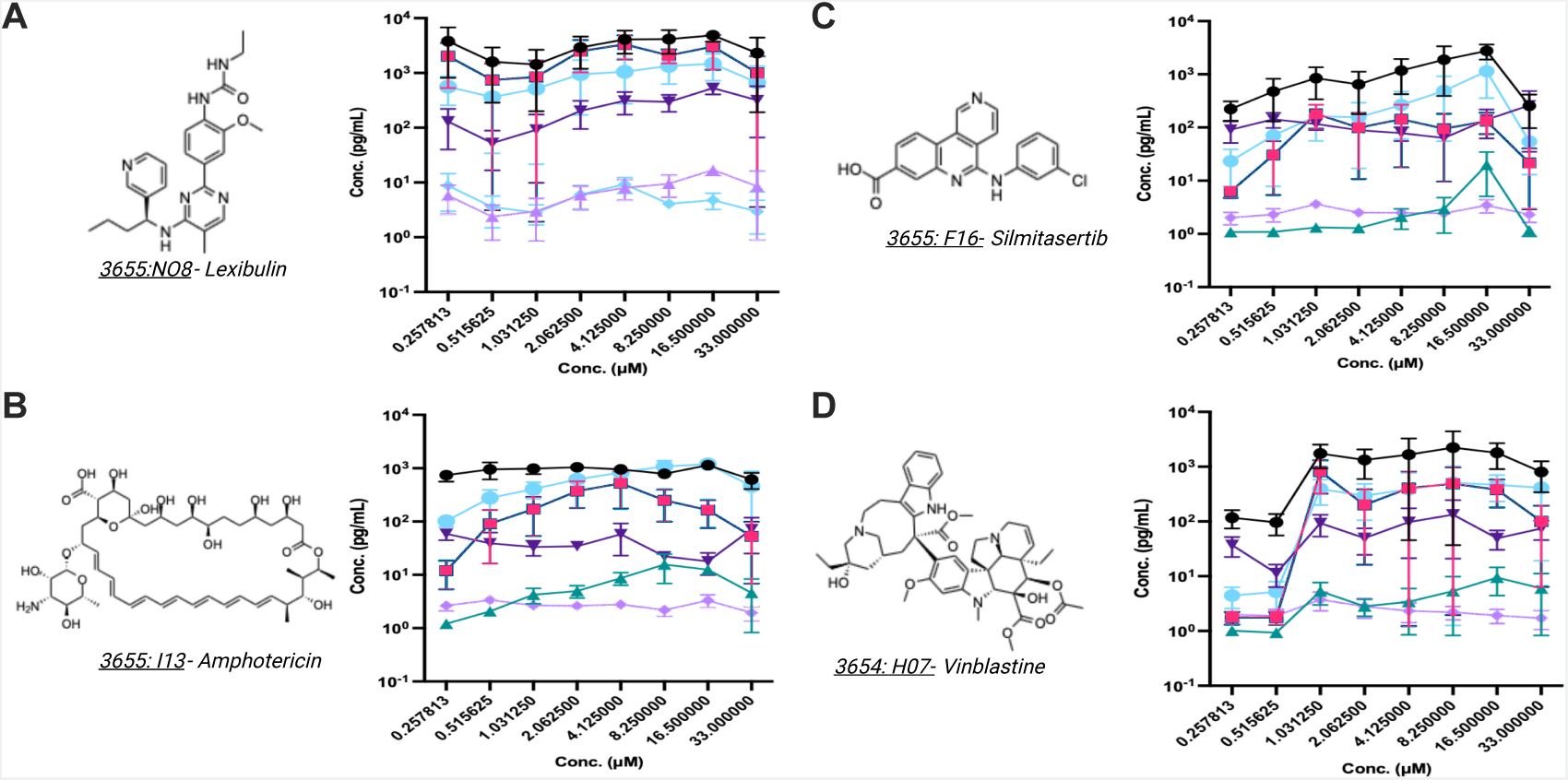
Experimental and compositional characteristics of the top four high throughput screen hits. Selection criteria for the top screening hits include demonstrations of titratability, robust efficacy in inducing multiple cytokines, and satisfactory potency. Chemical structure, well identification, and multiplexed concentration titration experimental data shown for these top screening hits: **A.** Lexibulin**, B.** Amphotericin, **C.** Silmitasertib, and **D.** Vinblastine.

Of note, one compound, not included in this list of four lead adjuvant candidates, triciribine phosphate, demonstrated very high efficacy in parts of the initial screen but did not perform well in downselection assays. Triciribine had significantly greater activity in nearly all of the AlphaLISA assays when compared to the next highest compound, demonstrating ∼thousand-fold greater activity in the confirmation experiment (Supplementary Figure 2). However, it had little activity in the multiplexed dose titration assays, including TNF induction (Supplementary Figure 3).

To investigate the source of this discrepancy, we tested triciribine with the other four adjuvant candidates in a quality assurance (QA) assay. In typical AlphaLISA assays, donor beads specifically bind to the cytokine of interest and acceptor beads attach to the same cytokine at a different site. When both beads come into close proximity to one another, a light wave is emitted, enabling measurements of overall luminescence to capture the concentration of the cytokine (Figure 6A). In this QA assay, compounds were tested in varying sets of conditions with respect to the assay reagents: with only the acceptor beads (Figure 6B), without any beads (i.e., molecules only) (Figure 6C), with only the donor beads (Figure 6D), or with both sets of beads (Figure 6E).

**Figure 6:**
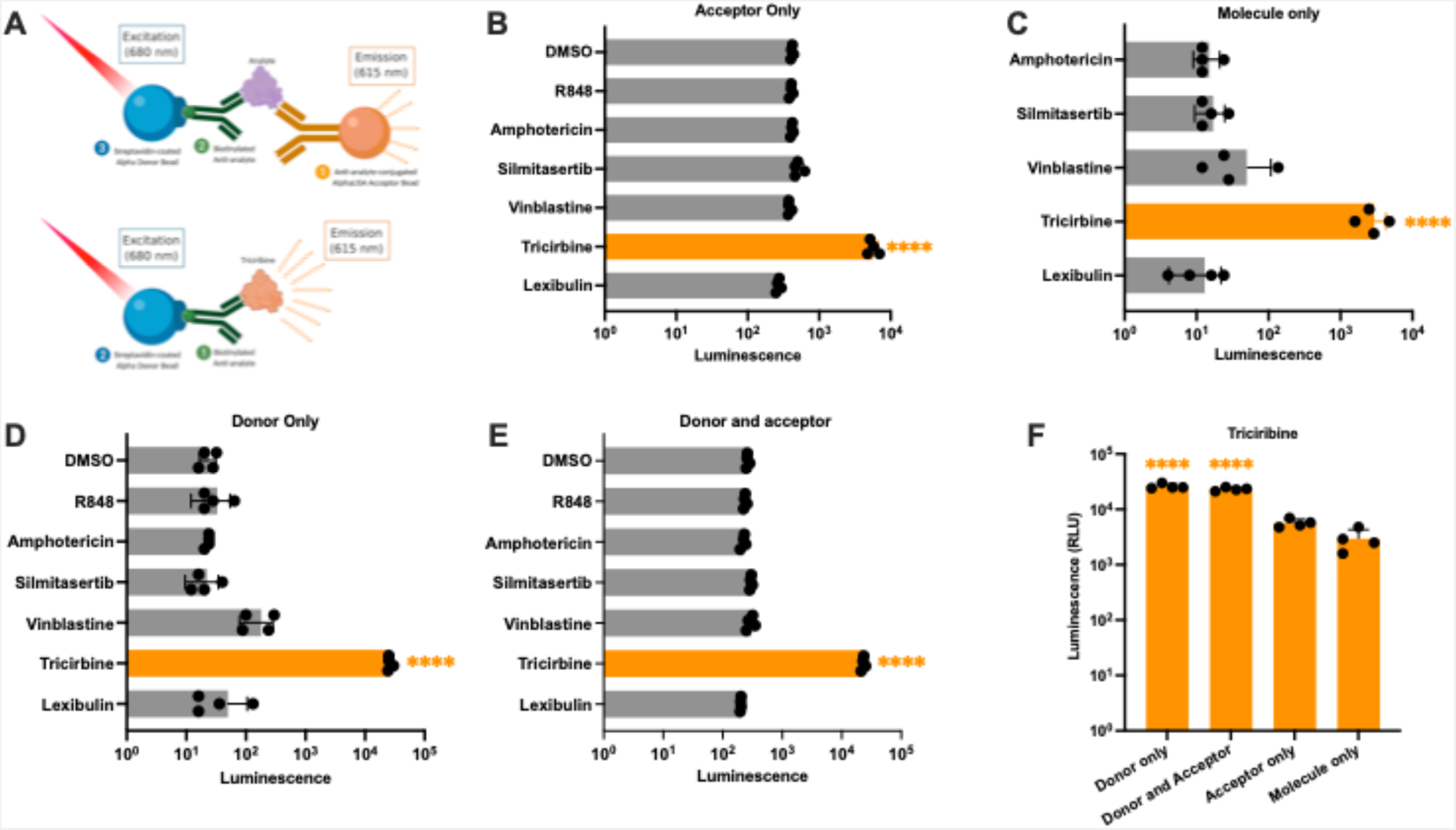
Quality assurance assay identifies a false positive compound that is an acceptor bead mimetic. Triciribine, a screening hit with low efficacy, low potency, and non-titratability, was found to be a potential acceptor compound in the AlphaLISA assay**. A**. Schematic representation of triciribine’s acceptor bead mimetic functionality.**B-E**. Luminescence of leading screen hits under different combinations of donor and acceptor bead additions. Results for each combination were analyzed by one-way ANOVA followed by post-hoc Tukey’s test for multiple comparisons. For statistical significance calculations: \**p* < 0.05 ** *p* < 0.01, **** *p* < 0.001,**** *p* < 0.0001**. F.** Luminescence of triciribine, under four AlphaLISA kit conditions summarized. Data for this compound were analyzed by one-way ANOVA followed by post-hoc Tukey’s test for multiple comparison

In all conditions, triciribine demonstrated a significantly larger luminescence (*p* < 0.0001, one-way ANOVA) when compared to all other tested compounds (Figure 6B-E). In fact, even without any reagent materials from the AlphaLISA assay, triciribine demonstrated appreciable luminescence (Figure 6C). This result suggests that triciribine likely functioned as an acceptor bead mimetic such that when it is excited with 680 nm light wave, the compound emits a 615 nm light wave that is picked up by the plate reader as an artificial signal (Figure 6D). As a result, triciribine represents a false positive for the cytokine measurements, as shown by this quality assurance assay. No similar trends were seen with other compounds, which further corroborated the adjuvant potential of these compounds.

### Post-HTS Evaluation of Adjuvantation

To further advance and validate these screening results in an accepted assay system, we tested the four adjuvant candidates as well as triciribine for TNF induction using a sandwich enzyme-linked immunosorbent assay (ELISA). As expected, the four adjuvant candidates demonstrated a sufficient TNF induction at a therapeutically relevant dosage of 11 µM (Figure 7A) and clear titratability (Figure 7B), thereby confirming the results of the previous rounds of screening. This result justified the examination of these compounds for potential adjuvanticity *in vivo*.

**Figure 7:**
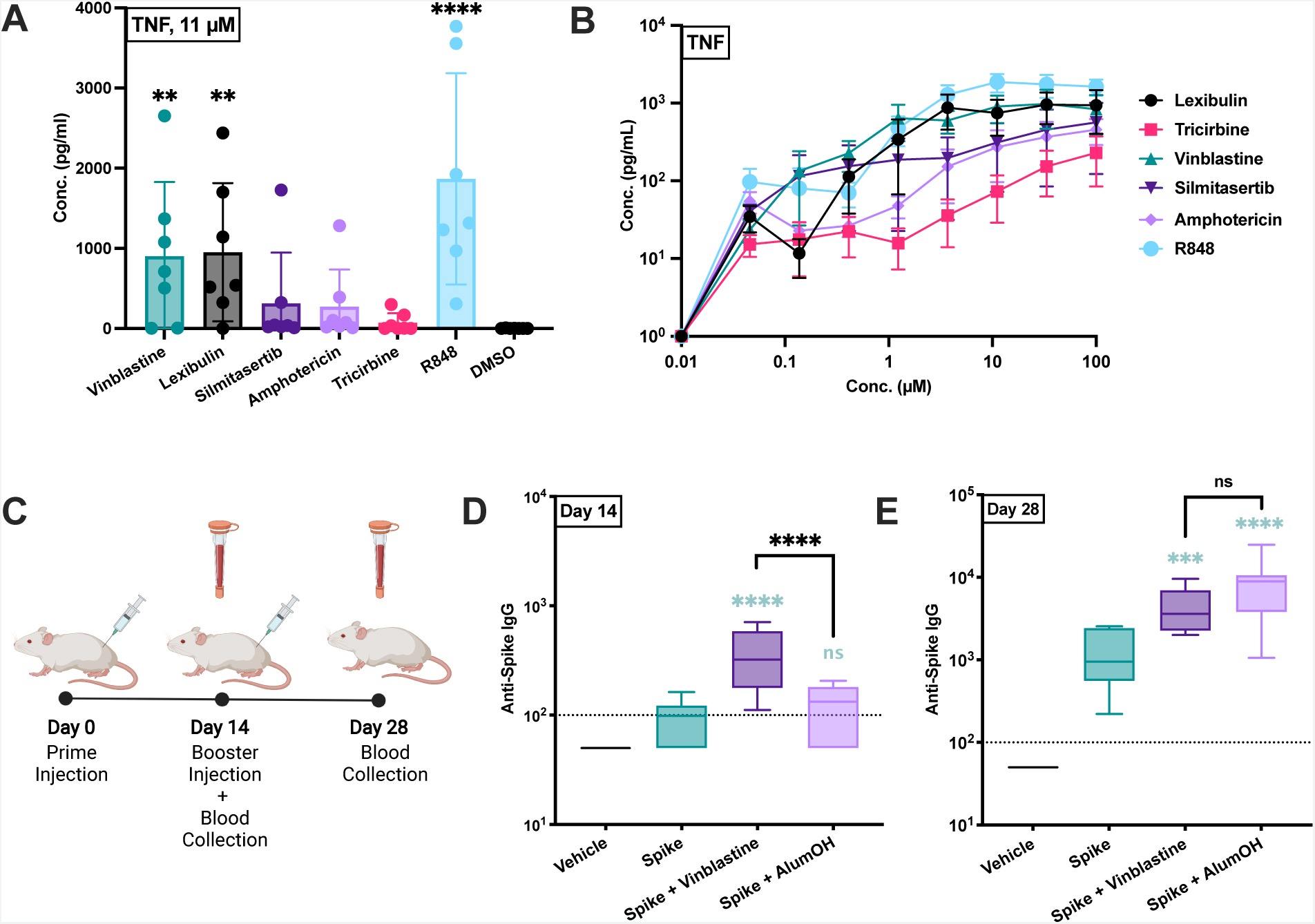
*Assessment* of downselected high-throughput hits for *in vitro* and *in vivo* adjuvanticity. **A**. TNF induction of leading screen hits at 11 µM. Data for each combination were analyzed by one-way ANOVA followed by post-hoc Dunn’s test for multiple comparison. Black stars indicate comparison to the DMSO control group. N=10 per condition. **B**. TNF induction by the four screening hits and triciribine phosphate for cryopreserved human PBMCs in a dosage-dependent manner. Results shown are mean ± SEM TNF concentration and fold change over DMSO negative control. N=10 per condition. **C.** Schematic representation of the prime-boost vaccination paradigm. Female 3-month-old BALB/c mice were immunized IM on days 0 and 14 with 1 µg of SARS-CoV-2 spike trimer. **D.** Anti-spike IgG titers at Day 14. Data for each combination were analyzed by one-way ANOVA followed by post-hoc Turkey’s test for multiple comparisons. Blue-green stars indicate comparison to the spike-only control group. N=7 per condition. **E.** Anti-spike IgG at Day 28. Data for each combination were analyzed by one-way ANOVA followed by post-hoc Turkey’s test for multiple comparison. Blue-green stars indicate comparison to the spike-only control group. N=7 per condition. For statistical significance calculations: \**p* < 0.05 ** *p* < 0.01, *** *p* < 0.001,**** *p* < 0.0001

We chose the top compound that demonstrated an appropriate efficacy and potency, vinblastine sulfate, for *in vivo* investigation. This compound was formulated with spike protein and administered in a 14-day prime-boost regimen (Figure 7C). At day 14, the vinblastine-adjuvanted group elicited significantly higher levels of anti-spike IgG than the spike-alone and spike-alum vaccine conditions [geometric mean titers (GMTs) of 308, 85, and 101, respectively; p < 0.0001] (Figure 7D). By Day 28, both the vinblastine-adjuvanted group (3992 GMT) and the spike-alum formulated group (6815 GMT) demonstrated greater anti-spike Ab titers than the spike-alone group (938 GMT). Of note, when compared to the spike-alum formulated group, the vinblastine-adjuvanted group demonstrated superior Ab titer production at Day 14 (p<0.0001) and non-inferior Ab titers at Day 28, (Figure 7E).

## Discussion

Novel vaccine formulations are some of the most impactful and effective strategies against the emergence and spread of infectious diseases, as illustrated by the current SARS-CoV-2 pandemic. However, current vaccine development pipelines cost an estimated $1–2 billion per formulation and can take decades to reach FDA approval (Light et al., 2009; Oyston and Robinson, 2012). Adjuvant discovery and formulation contribute to this financial and temporal bottleneck, largely due to the difficulty in finding sufficiently immunogenic, safe, and durable components that can improve vaccine efficacy. Phenotypic screens, which probe biologically relevant systems to discover novel phenotypes or functionality of compounds within a tested library, can aid this expensive and difficult discovery process (Moffat et al., 2017; Zheng et al., 2013). However, there have been very few successful phenotypic screening campaigns for the discovery of immunomodulatory compounds, much less for adjuvant discovery. Further, the few reported HTS campaigns incorporate single, well defined cell lines, limiting the biological relevance of their results. Here, we discuss a novel methodology and screening campaign that successfully leverages human primary cells to enable the identification of four screening hits demonstrating immunomodulation, with one compound presenting adjuvanticity. This methodology proffers a relatively cheap, fast, and effective strategy for adjuvant discovery.

Our screen identified four screening hits (Supplementary Table 3) that are known medications and bioactives used in a non-vaccine context. Of these, lexibulin, triciribine, and silmitasertib have known antiviral properties (Bouhaddou et al., 2020; Kalkeri et al., 2020; Porcari et al., 2003). Consistient with our findings, another hit, amphotericin, has been proposed as a TLR2 agonist adjuvant (Salyer et al., 2016). Our screen also identified our lead screening hit vinblastine sulfate, a vinka alkaloid extracted from the flowering herb *Vinca minor* (periwinkle) that prevents cell division by binding microtubular proteins in the mitotic spindle. While typically conceived of as an inhibitor of mitosis, vinblastine sulfate has also been noted to enhance DC maturation (Tanaka et al., 2009), consistent with our findings.

While phenotypic screens have been proposed to expand the pipeline of immunostimulatory adjuvants (Tom et al., 2019), few successful screens have been reported (Buckner et al., 2006; Garcia-Cordero et al., 2013; Wong *et al*., 2015). Further, these existing adjuvant-oriented screens exclusively utilize single, well defined cell lines, such as murine macrophage reporter cell lines or cultured bone marrow-derived dendritic cells. This approach enables the study of precise readouts and conditions in a reproducible and readily scalable manner, which is attractive considering the high levels of variability inherent in many screening paradigms (Ding et al., 2017; Fallahi-Sichani et al., 2013). However, the major limitation of this approach is that it fails to represent the robust and diverse functionality of the immune system, instead de-contextualizing cellular functions from the nuanced and highly interactive systems that exist in natural immunity.

Using human primary cells, as opposed to these conventional and non-diverse samples, enables the study of a variety of cell types and therefore provides a more robust and complete immunological profile of screened compounds. Notably, PBMCs include a diverse range of human immune cells including monocytes, dendritic cells, macrophages, T cells, B cells, and natural killer cells (Kleiveland, 2015). This cell composition captures not only the adaptive immune response from T and B cells also the innate immune response mounted by monocytes, NK cells, and importantly DCs. Monocytes, macrophages, and DCs are largely responsible for TLR-mediated responses in immunological contexts (Kawasaki and Kawai, 2014). However, TLR expression patterns have been demonstrated to be largely variable, including amongst these cell subtypes (Applequist et al., 2002; Zarember and Godowski, 2002). Thus, PBMCs can be valuable models of investigation in screening paradigms due to their capacity to advantageously capture broad TLR activity (Slavik *et al*., 1999) while also capturing the adaptive immune response mediated by innate immune activation. In turn, this approach may therefore enable more relevant and effective identification of adjuvant candidates in screening campaigns.

In our HTS platform, we tested 2,296 compounds, identified 197 hits, confirmed and downselected to 20 compounds, finalized four robust immunostimulatory leads, and tested the top candidate for *in vivo* adjuvanticity. When formulated with spike protein and tested in a murine *in vivo* model, the adjuvant candidate, vinblastine sulfate, enhanced humoral immune responses, as demonstrated by increased Ab titers compared to spike protein alone. Remarkably, this titer was significantly greater than the anti-spike Ab titers induced by an alum and spike formulation at Day 14 and was non-inferior to the alum-adjuvanted formulation at Day 28. Thus, starting from ∼2,500 compounds, this HTS program identified a compound, vinblastine sulfate, that performed comparably *in vivo* to the AH adjuvant benchmark. Of note, despite using human primary cells in a context of high-content HTS, the results were highly consistent, suggesting that our approach was able to control the added biological variability of utilizing primary cells while enabling the discovery of small molecule adjuvants.

Overall, our innovative screening approach presents clear advantages and strengths, including: (a) providing a rapid, reliable, and cost-effective system capable of testing immunomodulatory profiles of thousands of small molecules; (b) using an unbiased human primary immune cell screening platform, including the use of autologous plasma, a rich source of immunomodulatory factors (Pettengill *et al*., 2014; Sanchez-Schmitz et al., 2020; van Haren et al., 2016a), to enable more informative HTS evaluations as compared to more commonly leveraged cell lines; and (c) demonstrating system efficacy through the identification of three novel adjuvant candidates. Limitations of this methodology include: (a) inherent variability of human primary cells as compared to cell lines due to the diversity of human immune responses; (b) limited diversity of immunological readouts; and (c) utilization of chemical plates of known bioactive compounds, which have higher baseline likelihood of immunological activity. Neverthless, this methodology establishes the validity, potential, and precedent for larger-scale screens using human primary cells for adjuvant discovery.

Our screening methodology provides considerable flexibility in measuring soluble (eg, cytokine) and cell-associated (e.g., surface receptor) biomarkers. This powerful and innovative approach enables future screens that tailor soluble and cellular biomarkers for optimized discovery and precise functionality without sacrificing the added depth of information that comes from testing human primary cells. In future screens, more diverse and novel chemical libraries can be tested to discover truly novel small molecule adjuvants and further develop the existing library of approved and available adjuvants (Pulendran *et al*., 2021). These opportunities will hopefully allow researchers to push adjuvant discovery and formulation development to progress in parallel with our growing knowledge of prevalent infectious diseases.

## Stars Methods

### Key resource table

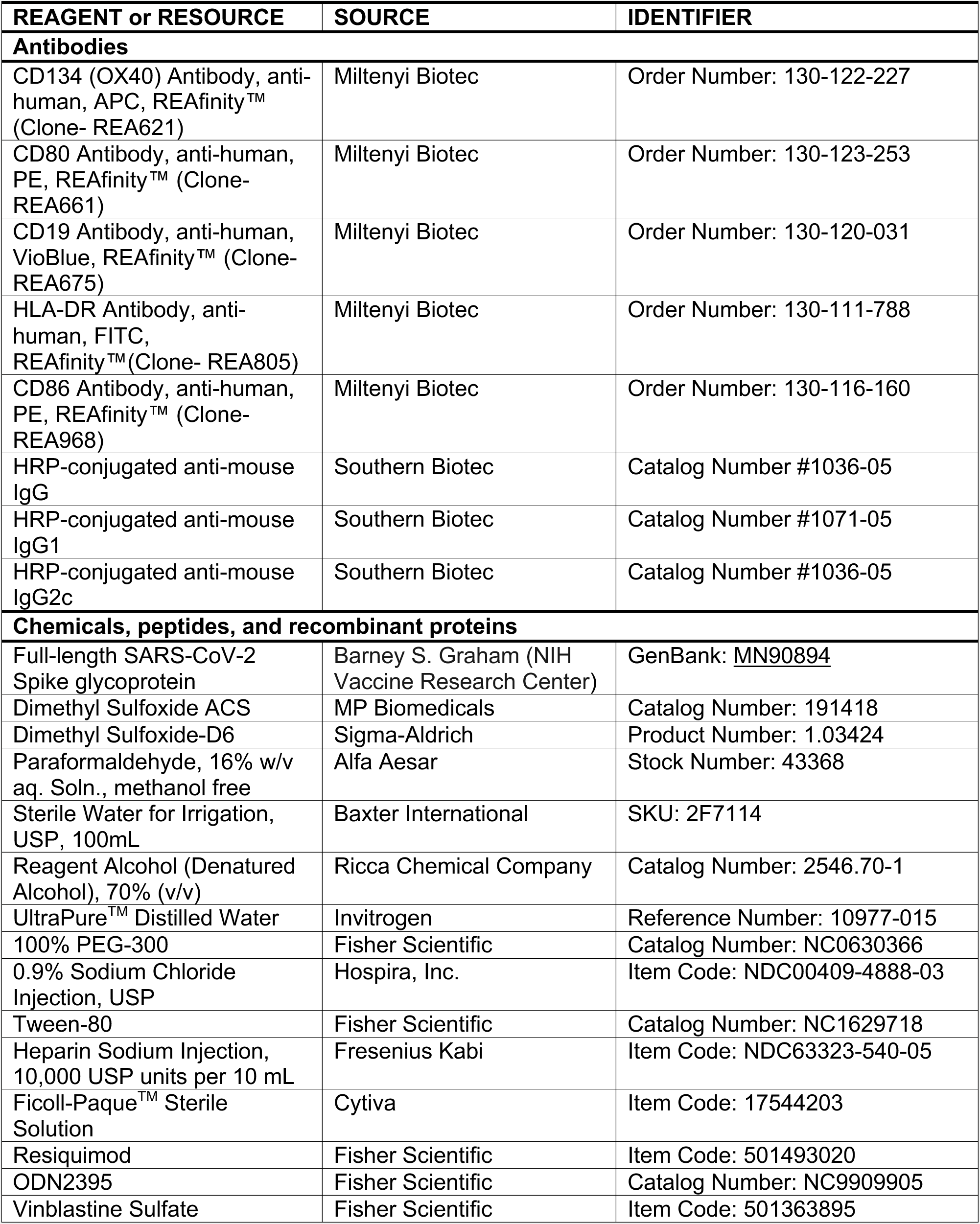
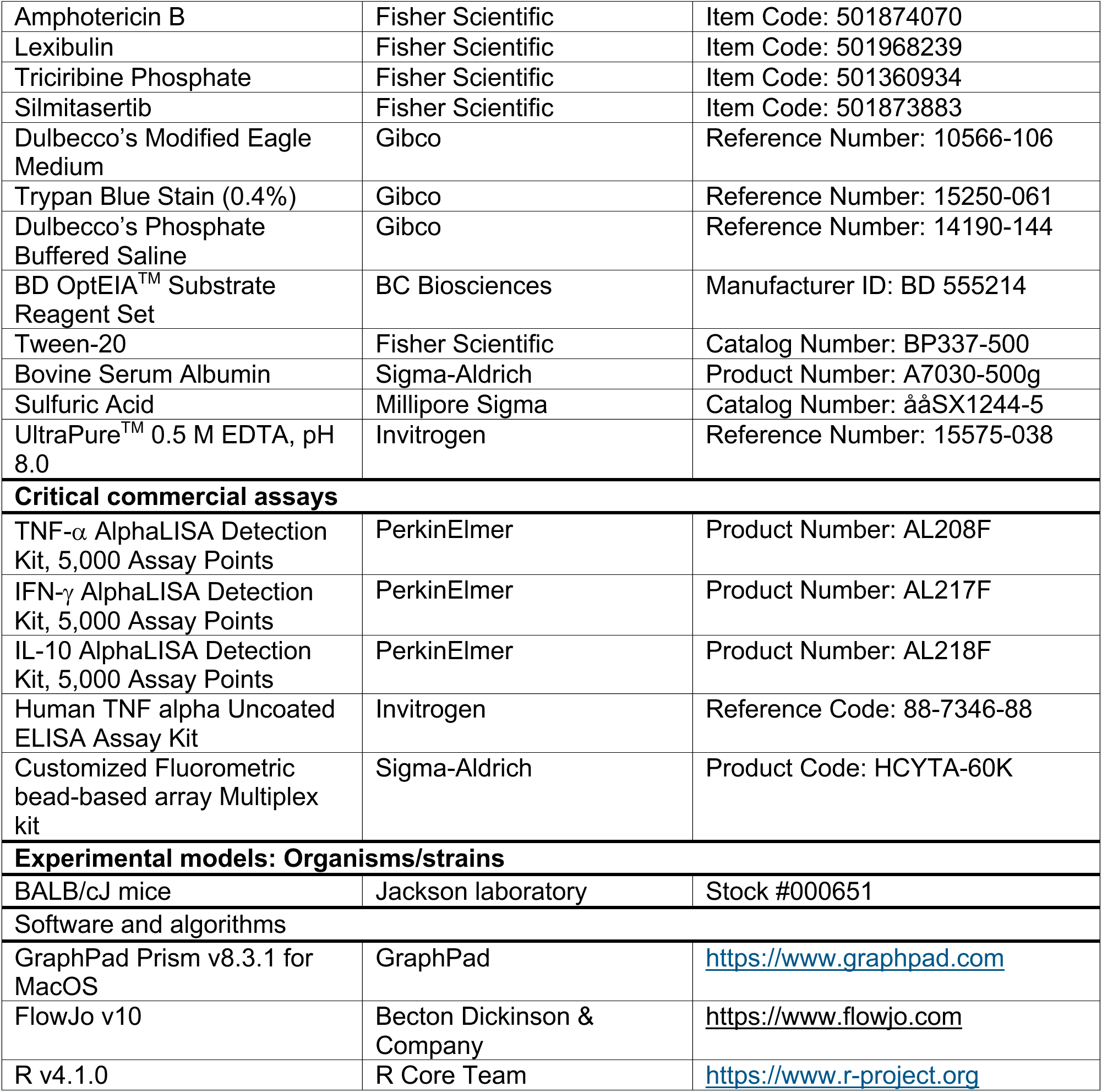

## Resource Availability

### Lead Contact

Further information and requests for resources and reagents should be directed to and will be fulfilled by the Lead Contact, Dr. David Dowling.

### Materials availability

This study did not generate new unique reagents.

### Data and Code Availabilåity

- All data reported in this paper will be shared by the lead contact upon request.
- Any additional information required to reanalyze the data reported in this article is available from the lead contact upon request.

## Experimental Model and Subject Details

### Human Peripheral Blood Samples

Peripheral blood (PB) was collected from healthy adult volunteers (n = 4 individual participants; Age and Sex: 23F, 25M, 27F, 27F) with approval from the Institutional Review Board of the Boston Children’s Hospital (protocol number X07-05-0223). Blood was anticoagulated with 1000 United States Pharmacopeia (USP) units heparin per mL of blood (Fresenius Kabi, Bad Homburg, Germany). Heparinized human blood was layered onto Ficoll-Hypaque gradients and the mononuclear cell (MC) layer was collected. PBMCs were washed 2 times with phosphate buffered saline (PBS), and live cell count determined using trypan blue (Sigma Aldrich; St Louis, MO). PBMCs were stored at -80°C with 50 million cells/1 mL for a total 1 mL per cryopreservation tube, with the liquid phase constituting 90% autologous plasma and 10% dimethyl sulfoxide (DMSO). After overnight freezing at 80°C in isopropanol-insulated containers to ensure linear freezing, the tubes were moved to long term liquid nitrogen storage. All study participants signed an informed consent form prior to enrollment.

### Animals

3-month-old female BALB/c mice were purchased from Jackson Laboratory (Bar Harbor, ME). Mice were housed under specific pathogen-free conditions at Boston Children’s Hospital, and all the procedures were approved under the Institutional Animal Care and Use Committee (IACUC) and operated under the supervision of the Department of Animal Resources at Children’s Hospital (ARCH) (Protocol number 19-02-3897R).

## Method Details

### Human PBMC Preparation for Screening

Cryopreserved human PBMCs were thawed for screening use. First, autologous plasma was thawed and centrifuged (25°C, 3000 x g, 10 min). Cryopreserved cells were thawed in a 37°C water bath for 3 min. 1 mL of autologous heparinized plasma was added to the tubes of cryopreserved cells and the cells were transferred to a 50 mL conical tube. Then, 3 mL of autologous plasma and 45 mL of DMEM were added dropwise as the tube was swirled. The cells were centrifuged at 500 g for 10 min and resuspended to 10 mL in DMEM with 10% autologous plasma, stained with trypan blue (Sigma Aldrich), and counted using a hemocytometer. Once counted, PBMCs were diluted in DMEM supplemented with 10% autologous plasma to a concentration of 50,000 cells/30 µL (1.67 x 10^6^ cells/mL). PBMCs in 10% autologous plasma were then added to 384 well Corning 3656 microplates (Corning, Corning, NY) using a Multidrop Combi Reagent Dispenser (ThermoScientific, Waltham, MA).

### Compound Libraries and Pinning

2,296 molecules were screened (Supplementary Table 1) and derived from the rolling and curated ICCB-Longwood Screening Facility at Harvard Medical School. All plates used a 384 well format. 704 molecules were chosen for known activity toward PBMCs; these molecules filled up two library plates and originated from the ChemDiv6 library (ChemDiv Inc, San Diego, CA). Another 1592 molecules were chosen from the Selleck bioactive chemical plates (Selleck Chemicals LLC, Houston, TX). The compounds (stored at 10 mM in dessicated conditions) were pinned at a volume of 100 nL, for a final concentration of 33 µM in a 384 well format. 5 µL of 0.3% DMSO(Millipore, Burlington, MA) diluted in DMEM, the negative control, and the TLR9 agonist ODN2395 and the TLR7/8 agonist R848 (both from Invivogen, San Diego, CA), the positive controls, were added to the wells manually for final concentrations of 1µM and 25 µM respectively. PBMCs were then stimulated for 72 hours at 37°C, 5% CO_2_ in a humidified ThermoScientific Forma CO_2_ Incubator (Waltham, MA).

### Supernatant Collection

After 72 hours, plates were centrifuged at 500 g for 10 min at 25°C with low brake, and supernatants collected using an Agilent Velocity 11 VPrep. 15 µL of supernatant were removed and transferred to an Eppendorf twin.tec PCR Plate 384 (Cat. 951020729, Hamburg, Germany). Plates were centrifuged at 500 g for 10 seconds and three aliquots of 2 µL of supernatant transferred to PerkinElmer AlphaPlate 6005359 (Waltham, MA) twice (using up a total of 4 µl of the supernatant). The three ΑlphaPlates were sealed with aluminum plate sealers and stored at - 80°C.

### Preparation for Flow Cytometry

After supernatants were harvested, plates with cells were centrifuged at 500 g for 10 min at 4°C and the remaining supernatant removed. Using a Combi reagent dispenser, 20 µL of cold 250 mM EDTA was added to each well. Plates were shaken using a Labline 4625 Titer Plate Shaker (Labline Instruments, Melrose Park, IL) at 700 rpm for 10 min and then centrifuged at 750 g for 10 minutes at 4°C. EDTA was removed and cells were washed with 30 µL of PBS. After the wash, an antibody cocktail consisting of Miltenyi REAfinity Recombinant antibodies HLA-DR Ox40 APC REA621 (Bergisch Gladbach, Germany) was diluted 1:100 in PBS and 5 µL were added to each well using the Combi reagent dispenser. Plates were incubated at 4°C for 1 hr prior to washing cells with 40 µL PBS. Plates were centrifuged at 750 g and 20 µL of 1% paraformaldehyde was added to each well. Plates were then stored at 4°C. Before acquisition of cells by flow cytometry (which occurred at least one day but no longer than a week post fixation), plates were centrifuged at 750 g for 10 min at 4°C, paraformaldehyde removed via a wash and 20 µL of PBS was added.

### Flow Cytometry on iQue

High-throughput flow cytometry was completed using a Sartorius IntelliCyt iQue3 with 384 well cellular acquisition, employing iQue Forecyt Software (Essen BioScience, Ann Harbor, MI). Acquisition was designed to analyze populations of monocytes, B cells, and T cells, with the following percent positive metrics calculated for each well: HLA-DR^+^ Monocytes, CD80/CD86^+^ Monocytes, HLA-DR/CD80/CD86^+^ Monocytes, HLA-DR^+^ B Cells, CD80/CD86^+^ B Cells, HLA-DR/CD80/CD86^+^ B Cells, and Ox40^+^ T cells. Supplementary Figure 4 describes the typical gating strategy.

### Cytokine Quantification for the PBMC Screen

Concentrations of tumor necrosis factor alpha (TNF), interferon gamma (IFN-γ), and interleukin 10 (IL-10) were quantified in the supernatant from the PBMCs using an AlphaLISA assay (PerkinElmer, Waltham, MA). AlphaPlates containing 2 µL of supernatant were thawed at 25°C for 15 minutes. The provided assay procedure was followed, and the reagents were dispensed using a Combi reagent dispenser. In brief, for TNF, Anti-TNFα Acceptor beads and Biotinylated Antibody Anti-TNFα Mix were prepared per the manufacturer’s instructions and 8 µL of this cocktail were added to all wells. The plates were centrifuged at 500 g for 10 min at 25°C with low brake to bring the reagents to the well bottom. Plates were covered but not sealed and incubated in the dark at 25°C for 1 hr. Streptavadin Donor bead mixture was prepared per the manufacturer’s instructions and 10 µL were added to all wells. Plates were covered with another plate and incubated at 25°C in the dark for 30 minutes. For IFN-γ, the same procedure was followed except for the reagents in the respective kit. For IL-10, the mixes were prepared and 2 µL of the Anti-IL-10 Acceptor beads and Biotinylated Antibody Anti IL-10 Mix was added to each well. The plates were incubated for 60 min and then 16 µL of the Streptavadin Donor bead mixture was added to each plate. Plates were read using a PerkinElmer Envision Plate Reader (PerkinElmer, Waltham, MA) according to the following specifications: A1-384 aperture, Mirror Module Barcode 444, EMS filter Barcode 244 (570nm).

### Hit Calling Methodology

All well-based data from the high throughput AlphaLISA luminescence readings were exported as comma-separate values (CSV) files from the Envision Plate Reader. High throughput flow cytometry data were analyzed employing iQue ForeCyt analysis software and results exported as CSV files. Data were initially analyzed and transformed by the quality assurance team at ICCB-L. Statistical analyses were conducted on a plate-by-plate basis. A Z’ factor was calculated for each assay plate based on positive controls (25 µM R848 and 1 µM ODN2395) in column 24 and negative controls (0.3% DMSO) in column 23. Prior to hit calling for the luminescence readings from all AlphaLISA readouts, luminescent intensity values were log_10_-transformed and the median of each plate calculated. The absolute difference between the log_10_ of each value and the median of the corresponding plate was calculated. The median of the absolute differences was also calculated. From the median and median absolute difference, a z-score based on the log_10_ of each value was calculated. For hit calling criteria, if the z-score was >2 for both duplicates, the molecule was considered a hit for that donor in that readout. The hit calling methodology for the percent positive readings from the high-throughput flow cytometry was identical to the luminescence readings except that instead of log_10_, the raw percent positive was used in the calculation of the plate median, median absolute difference, and z-score. Each of the 2,296 compounds screened in the PBMC screen was tested in duplicate for each of the four donors. A molecule was registered as a potential hit for a particular donor if any of the readouts registered the molecule as a hit. Further, a molecule was moved onto the confirmation screen if it registered as a potential hit in two of the four donors.

### Confirmation Screen Readouts and Hit Calling

To more closely align with the results seen in the primary screen, PBMCs from the same four donors were utilized. PBMCs were thawed, cultured in 10% autologous plasma and 90% DMEM, and plated as in the primary screen. The 197 potential hit compounds were arrayed into a polypropylene 384-well at a concentration of 10 mM and a volume of 1.5 µL. 13.5 µL of DMEM was added to each well and 1 µL was then added to the plated PBMCs for a final concentration of 33 µM, as per the primary screen. PBMCs were incubated with the molecules for 72 hr. Harvesting of supernatants, cytokine assay, and flow cytometry were performed as in the primary screen. As the hit calling method in the primary screen could not be employed (z score calculated based on the entire plate would give inaccurate hits), a new method of z score calculation was used. A z-score was determined for each value based on the average and standard deviation of the negative control (DMSO). Accordingly, thresholds were adjusted so that a z-score > 3 in both duplicates was a hit in that donor for a given readout. If a compound scored positive in one donor for cytokine or monocyte readouts, regardless of whether this was the positive readout from the primary screen, it was considered a confirmed hit and advanced to a dose titration confirmation.

### AlphaLISA Quality Assurance Assay

For quality assurance (QA), we assessed whether hit molecules truly induced cytokine production or were false positives by artefactually enhanced signal in the AlphaLISA assay. Confirmed compounds were incubated with Anti-TNF Acceptor beads and Biotinylated Antibody Anti TNF Mix and Streptavadin Donor beads, Anti-TNF Acceptor beads and Biotinylated Antibody Anti TNF Mix only, or Streptavadin Donor beads at concentrations of 33 or 100µM and incubated per the AlphaLISA protocol as above. For the conditions that only received one type of bead, buffer without beads was added at the other time point (eg. Acceptor only condition received 10 µL of buffer instead of donor beads). Plates were read using an Envision Plate Reader per the following specifications: A1-384 aperture, Mirror Module Barcode 444, EMS filter Barcode 244 (570nm).

### Dose Titration Confirmation

PBMCs from the same four donors were used in order to align with the results seen in the primary and confirmation screen. PBMCs were thawed and plated in the same manner as the primary and confirmation screen. Using a Hewlett Packard HPD300e, compounds were serially diluted in duplicate at 1:2 in an 8-point dose titration curve with a top concentration of 33µM and 0.3% DMSO in each well. Controls included 0.3% DMSO, 1 µM ODN 2395, and 25 µM R848. PBMCs (30µL, 50,000 per well) were then plated on top of the molecule using a Combi reagent dispenser. PBMCs were incubated with the molecules for 72 hr prior to harvesting supernatants using an Angilent Vprep. 10 µL of the supernatants were stored in Eppendorf twin.tec PCR Plates at -80°C and subsequently analyzed for IFN-γ, IL-1β, IL-10, IL-12p70, CXCL-10, and TNF using a fluorometric bead-based array Multiplex kit (Millipore; Billerica, MA) and a Luminex Multiplex Instrument (Millipore), following the manufacturer’s recommendations.

### Further *in vitro* and *in vivo* Assays

Cryopreserved PBMCs from seven additional study participants were used to evaluate the top hit molecules in further *in vitro* assays. PBMCs were thawed in the same manner as the primary and confirmation screens and diluted to 300,000 cells/180 µL (1.67 x 10^6^ cells/mL). The top hit molecules from the screen were run side-by-side with negative (1% DMSO) and positive (R848; top 100µM) controls in an 8-point dose-response experiment starting at a top concentration of 100 µM and going down to 46 nM, with each molecule and R848 tested in triplicate for each donor. DMSO was tested in duplicate per donor. 180 µL of cells and 20 µL of diluted compound were added to a Falcon 353227 TC-treated U-Bottom 96 well plate (Corning, Corning NY). The cells were then stimulated for 24 hours at 37°C, 5% CO_2_ in a humidified ThermoScientific Forma CO_2_ Incubator (Waltham, MA). After 24 hours, the plates were spun down at 500 g for 10 minutes at 25°C with low brake and the supernatants were manually collected. TNF concentrations in the supernatants were measured by ELISA (Fisher Scientific, Waltham, MA, Cat# 88-7346).

The top adjuvant candidate, Vinblastine, was injected into mice at 50 nmol per treatment. Mice were injected with 1 µg of full-length SARS-CoV-2 spike glycoprotein (M1-Q1208, GenBank MN90895) formulated with or without candidate adjuvants. A mock treatment group received a 50 µL injection of phosphate-buffered saline (PBS). Intramuscular injections were administered in the caudal thigh on days 0 and 14, and serum was collected 2 weeks following each immunization. Serum IgG, IgG1, and IgG2a concentrations were measured by antibody binding ELISA, using an established protocol.

## Quantification and Statistical Analysis

Statistical analyses employed Prism v9.0.2 (GraphPad Software) and R software environment v4.0.4. Data were analyzed by one-way ANOVAs followed by post-hoc Tukey’s test for multiple comparisons in Figure 7. For all statistical significance: \**p* < 0.05 ** *p* < 0.01, **** *p* < 0.001,**** *p* < 0.0001. Individual n values in figure 7A-B represents individual human samples from different study participants and individual n values in Figure 7D-E represent individual mice. Mean and SEM was used for all precision measures.

## Graphics

Figures 1, 3E, 6A, and 7C were made using Biorender.com.

## Acknowledgements

We thank the members of the BCH *Precision Vaccine Program* for helpful discussions. We thank Drs. Kevin Churchwell, Gary Fleisher, David Williams, and Mr. August Cervini for their support of the *Precision Vaccines Program*. D.J.D. thanks Ms. Siobhan McHugh, Ms. Geneva Boyer, Mrs. Lucy Conetta and the staff of Lucy’s Daycare, the staff the YMCA of Greater Boston, Bridging Independent Living Together (BILT), Inc., and the Boston Public Schools for childcare and educational support during the COVID-19 pandemic. This study was supported in part by US National Institutes of Health National Institutes of Allergy and Infectious Diseases (NIAID) awards, including Adjuvant Discovery (HHSN272201400052C and 75N93019C00044) and Development (HHSN272201800047C) Program Contracts to O.L. D.J.D.’s laboratory is supported by NIH grant (1R21AI137932-01A1), Adjuvant Discovery Program contract (75N93019C00044). The *Precision Vaccines Program* is supported in part by the BCH Department of Pediatrics and the Chief Scientific Office. The high throughput screen was conducted at the ICCB-Longwood Screening Facility at Harvard Medical School and the authors thank the ICCB-L team for their support.

## Author contributions

BL and KC designed, performed, and analyzed the experiments in addition to writing and editing the manuscript. EN, TRO, MDL, DS, performed the experiments and edited the manuscript. JAS provided design feedback and edited the manuscript. HSS and SDP expressed and purified SARS-CoV-2 spike protein. AO and JBS provided design feedback and contributed to the statistical analysis. OL, SVH, and DJD conceived the project, designed the experiments, and edited the manuscript.

## Supplementary Figures

**Supplementary Figure 1:**
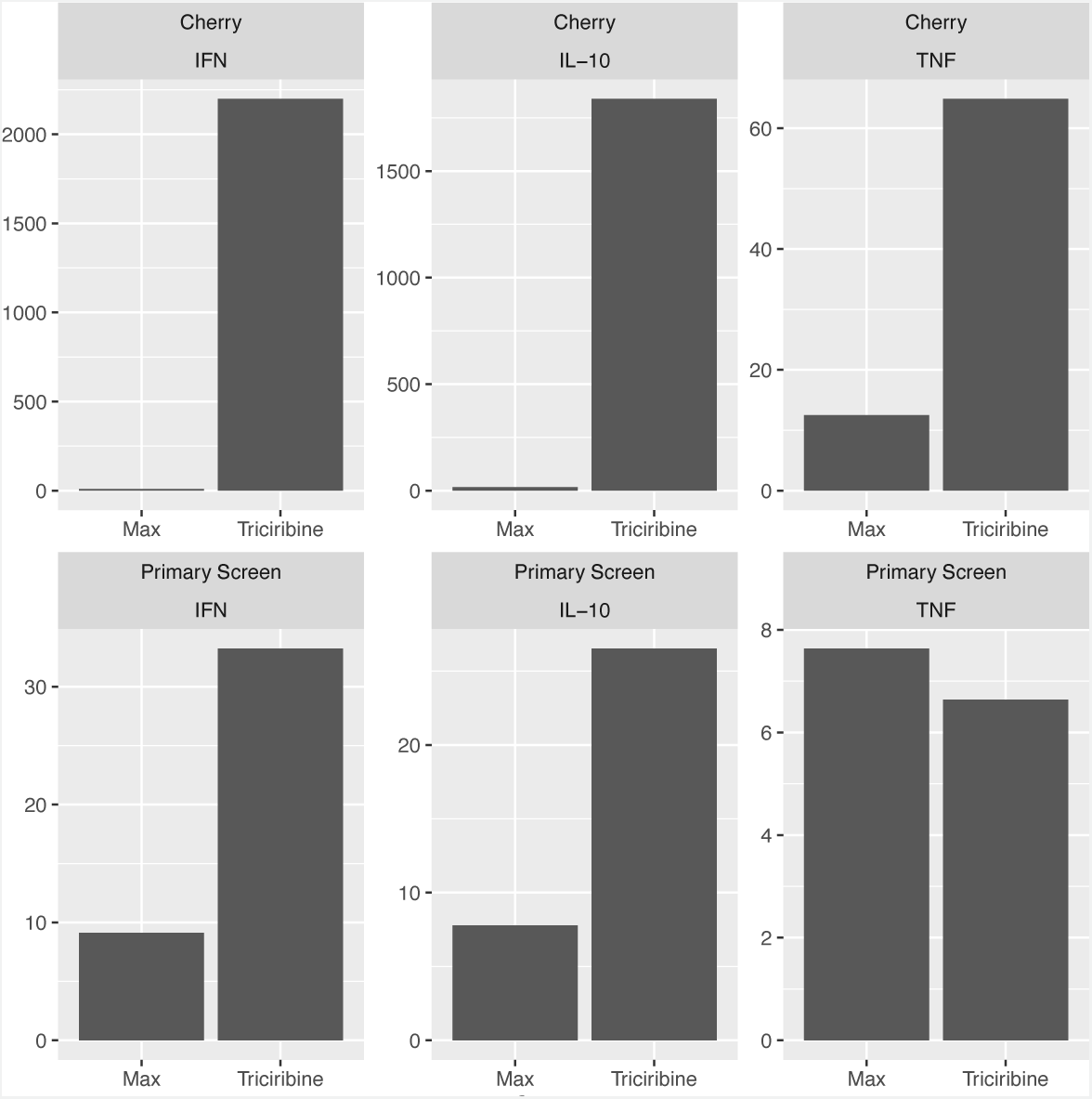
Triciribine Phosphate Historical AlphaLISA Performance. Triciribine average z score in each AlphaLISA assay is compared to the highest non-Triciribine z score in the corresponding experiment. While the raw z scores are calculated differentially in the cherry pick experiment and the primary screen, triciribine demonstrated a consistently sizeable luminescence that ranges from ∼3 fold up to ∼1000 fold greater than the next highest luminescence.

**Supplementary Figure 2:**
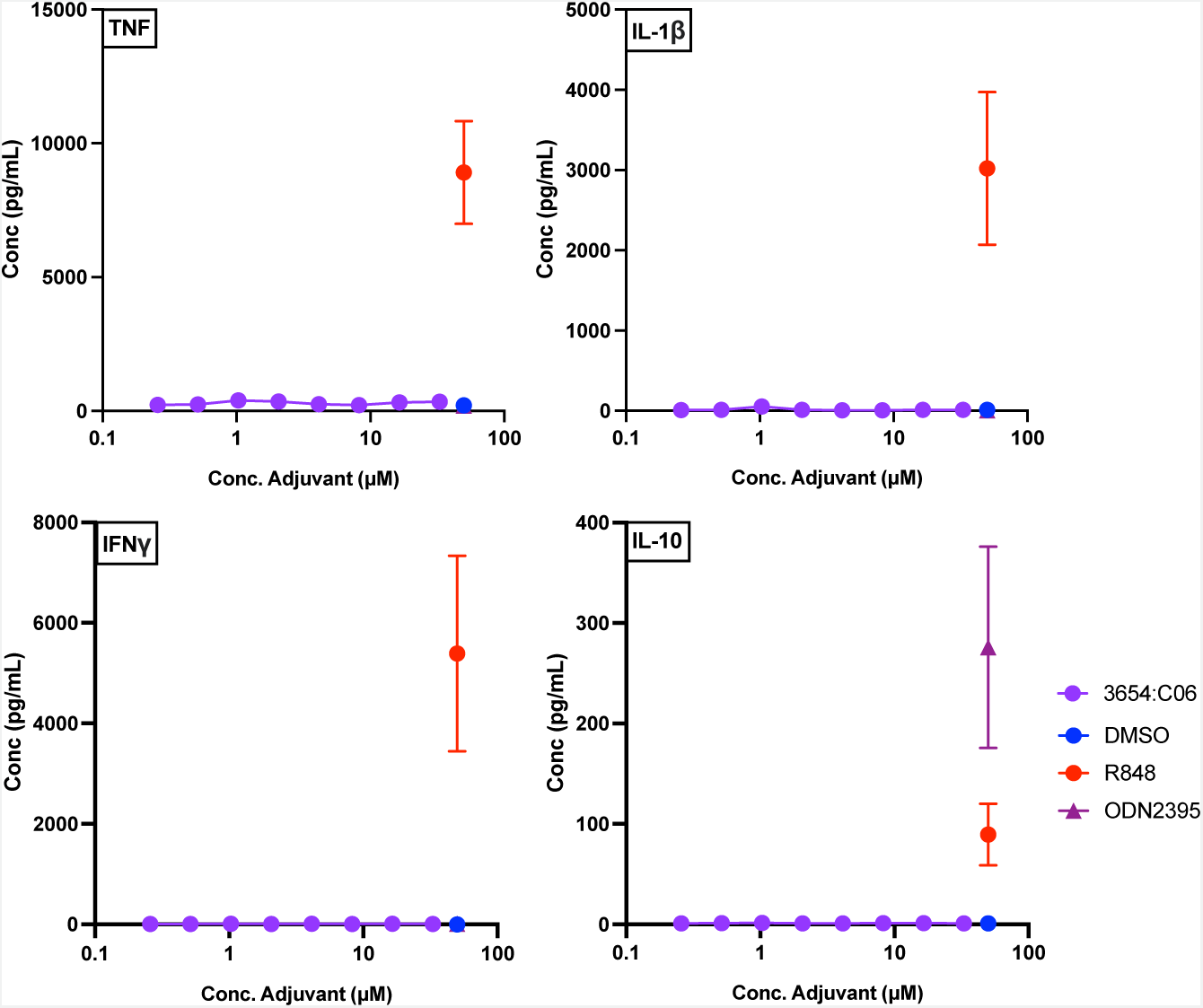
Triciribine Phosphate Concentration Titration Assay Results. Human PBMCs were cultured in 90% DMEM and 10% autologous PPP and stimulated with triciribine phosphate at an 8-point concentration titration from 0.25 µM to 33 µM. Cytokine induction was measured using a Millipore multiplex assay. Evidently, Triciribine demonstrates a weak cytokine induction outside of the AlphaLISA assay system, indicating that Triciribine is a weak immunomodulator.

**Supplementary Figure 3:**
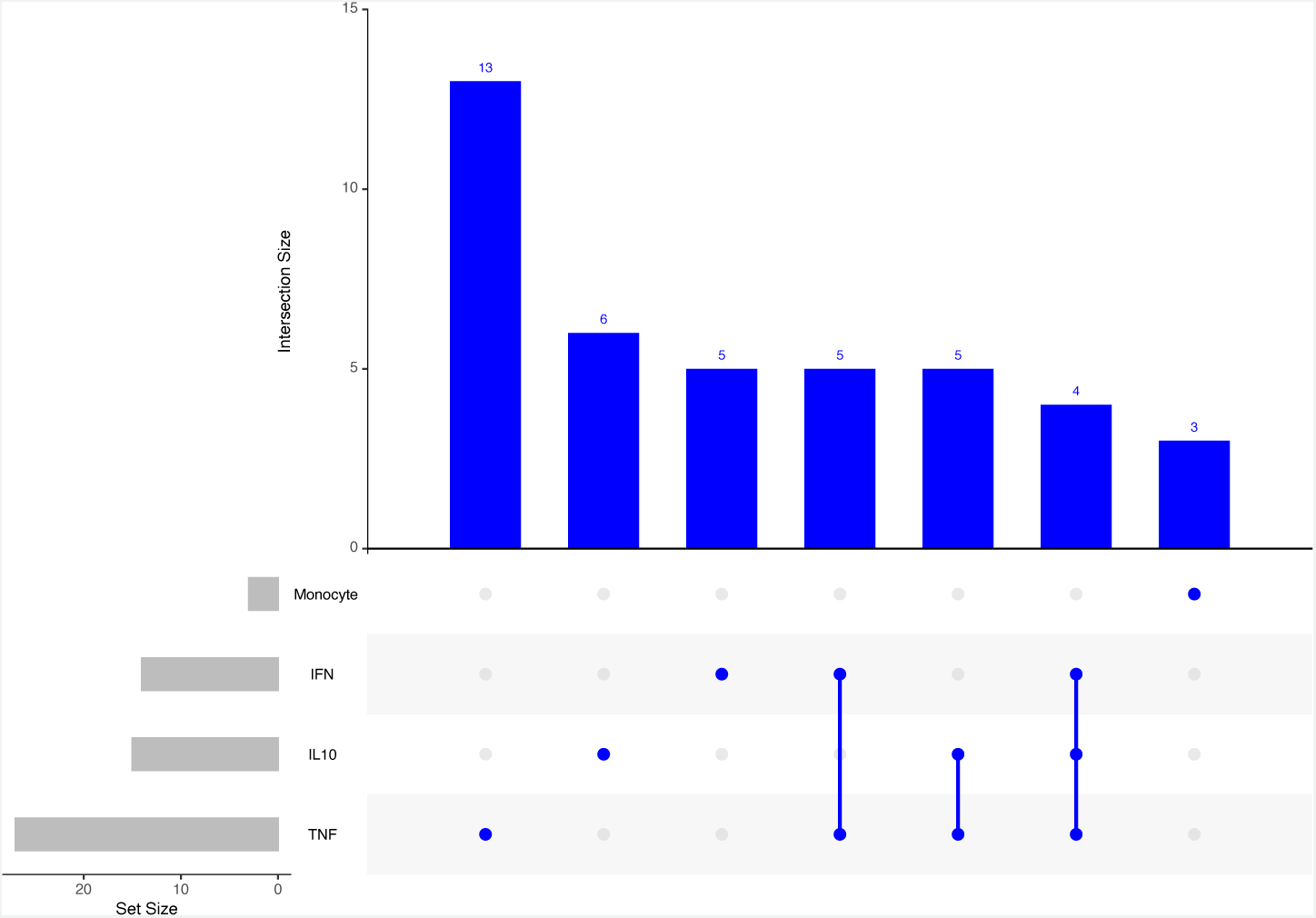
UpSet Plot demonstrating Intersections of Biomarkers from the Confirmation Assay. Frequencies of intersecting hit biomarkers demonstrate a non-uniform and non-predictive relationship between biomarker system.

**Supplementary Figure 4:**
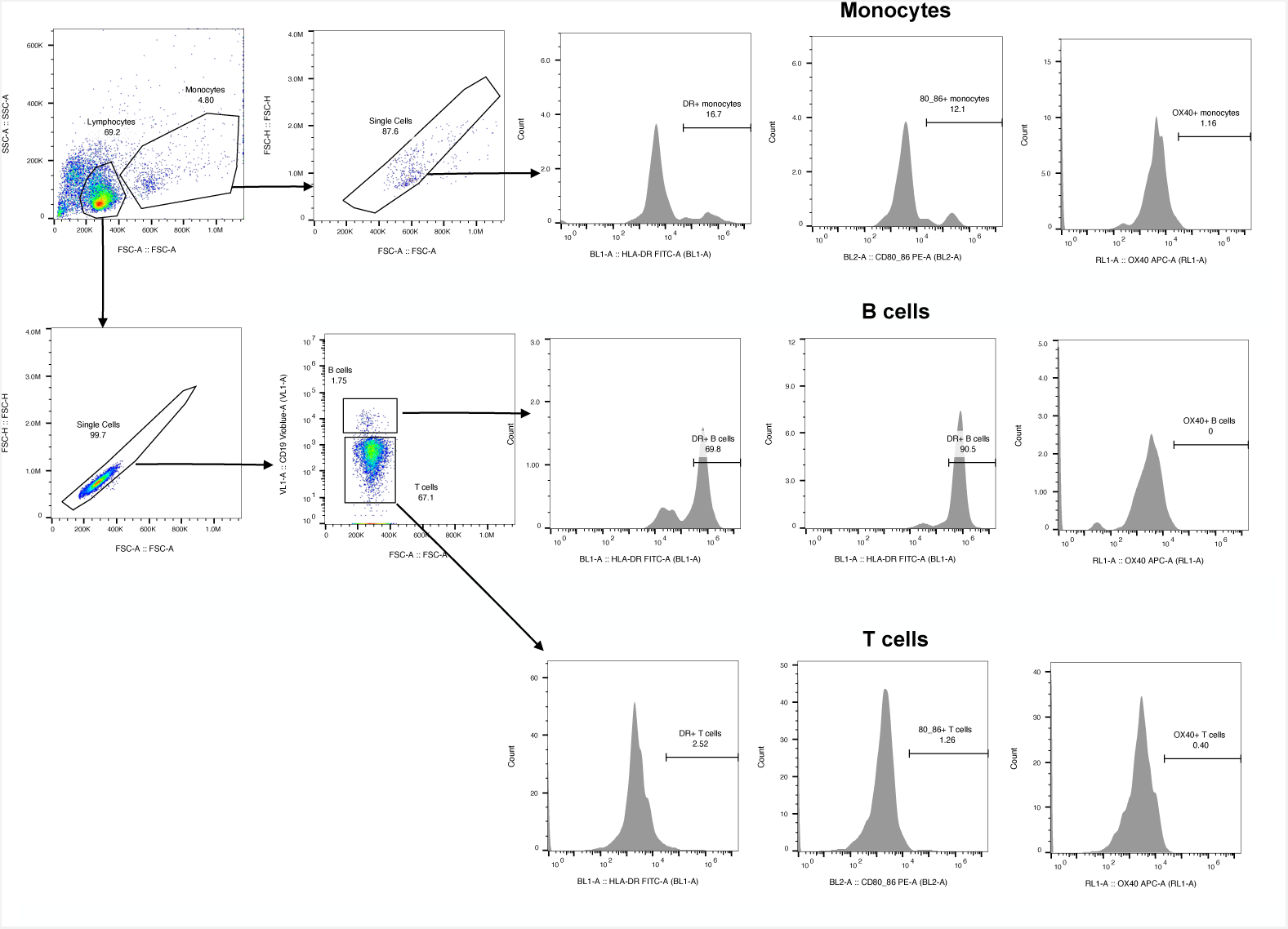
Representative Gating Strategy. The gating strategy was used for the identification of activated or naiive B cells, T cells, and monocytes. The activation markers used were HLA-DR and CD80/86 for monocytes and B cells as well as Ox40 for T cells.

**Supplementary Table 1: Screened Compounds**

The 2,296 compounds are identified by their compound name (if applicable), well and plate identification, molar concentration, and chemical vendor.

**Supplementary Table 2:** Potency and Efficacy Summaries for Screening Finalists in the Concentration Titration Experiment. Potency and efficacy as measured by EC_50_ and maximal cytokine induction for each biomarker shown for **A.** Lexibulin**, B.** Amphotericin, **C.** Silmitasertib, and **D.** Vinblastine Four parameter line curve estimations and EC_50_ calculations employed GrahPad Prism 9.0.

| A | Cytokine | Log EC<sub>50</sub> | Log EC<sub>50</sub> Rank | Max Efficacy (pg/mL) | Max Efficacy Rank |
| --- | --- | --- | --- | --- | --- |
|  | TNF | 2.414 | 6 | 3867 | 1 |
|  | IL10 | 0.6176 | 9 | 12.23 | 2 |
|  | IP10 | 0.4567 | 12 | 398.2 | 5 |
|  | IFN | 0.1167 | 8 | 1897 | 3 |
|  | IL1β | 0.2333 | 11 | 1140 | 3 |
|  | IL12P70 | 1.295 | 14 | 5.871 | 2 |

| B | Cytokine | Log EC<sub>50</sub> | Log EC<sub>50</sub> Rank | Max Efficacy (pg/mL) | Max Efficacy Rank |
| --- | --- | --- | --- | --- | --- |
|  | TNF | 5.955 | 11 | 844.8 | 5 |
|  | IL10 | 0.3946 | 7 | 10.72 | 4 |
|  | IP10 | -0.9525 | 2 | 71.715 | 17 |
|  | IFN | 0.01218 | 6 | 271.6 | 6 |
|  | IL1β | 0.1478 | 10 | 905.6 | 4 |
|  | IL12P70 | 1.668 | 18 | 2.795 | 9 |

| C | Cytokine | Log EC<sub>50</sub> | Log EC<sub>50</sub> Rank | Max Efficacy (pg/mL) | Max Efficacy Rank |
| --- | --- | --- | --- | --- | --- |
|  | TNF | 3.959 | 8 | 1627 | 3 |
|  | IL10 | 0.9383 | 11 | 10.5 | 5 |
|  | IP10 | 1.235 | 16 | 260.1 | 8 |
|  | IFN | -0.2688 | 3 | 112 | 8 |
|  | IL1β | 0.6775 | 15 | 576.4 | 5 |
|  | IL12P70 | -0.2816 | 3 | 2.806 | 8 |

| D | Cytokine | Log EC<sub>50</sub> | Log EC<sub>50</sub> Rank | Max Efficacy (pg/mL) | Max Efficacy Rank |
| --- | --- | --- | --- | --- | --- |
|  | TNF | 0.0636 | 3 | 1622 | 3 |
|  | IL10 | 0.4163 | 8 | 9.382 | 6 |
|  | IP10 | -0.1818 | 8 | 82.49 | 16 |
|  | IFN | -0.2352 | 5 | 398.7 | 4 |
|  | IL1β | -0.08816 | 6 | 413.5 | 6 |
|  | IL12P70 | 0.9511 | 10 | 2.578 | 11 |

**Supplementary Table 3:** Top HTS Campaign Hits. The four lead screening hits are described by their utility as known bioactives in addition to their current regulatory status. Any relevant recorded immunological activity of these hits and known mechanisms of action relevant to their bioactive status are included.

| Drug | Diseases | Regulatory Status | Previously Recorded Immunological Activity | Alternative Mechanism of Action |
| --- | --- | --- | --- | --- |
| **Lexibulin** | Multiple Myeloma, Glioblastoma. | Phase II Discontinued | Potential Antiviral Activity | Rapid Reorganization of Microtubules |
| **Vinblastine** | Breast cancer, Testicular cancer, Neuroblastoma, Hodgkin's and Non-Hodgkin's Lymphoma, and Histiocytosis. | FDA Approved | Dendritic Cell maturation, Potential Adjuvanticity. | Microtubule Assembly Inhibition |
| **Silmitasertib** | Breast Cancer*, Severe Covid 19*, Advanced Cholangiocarcinoma**. | Phase II*, Orphan Drug Status** | Potential Antiviral, Antifungal Activity | Protein Kinase CK2 Inhibition |
| **Amphotericin B** | Fungal Infections, Visceral Leishmaniasis, Primary Amoebic Meningoencephalitis. | FDA Approved | Antifungal and Antiparasitic Activity | Cell Membrane Disruption |

